# Genomic epidemiology of emerging terbinafine-resistant Trichophyton indotineae

**DOI:** 10.1101/2024.10.29.620818

**Authors:** Johanna Rhodes, Sui Ting Hui, Sarah Dellière, Richard C. Summerbell, James A. Scott, Amtoj Kaur, Richard C. Barton, Rodrigo Leitao, Samuel Hemmings, Rebeca Goiriz, Jonathan Lambourne, Rhys A. Farrer, Silke Schelenz, Roderick J. Hay, Andrew M. Borman, Anuradha Chowdhary, Alireza Abdolrasouli, Matthew C. Fisher

## Abstract

Dermatophyte skin infections affect around a quarter of the world’s population and are a growing public health concern due to increasing incidence of novel species causing severe infections that are resistant to antifungal treatments. Trichophyton species cause the greatest burden of dermatophytosis worldwide, with the T. mentagrophytes species complex being particularly associated with the emergence of new aggressive infections. One emerging species, T. indotineae (originally T. mentagrophytes genotype VIII) is notable for the extensive nature of the often inflammatory infection, its clinical resistance to terbinafine antifungal treatment, and its rapid global spread. To better understand the epidemiology of this disease, we sourced isolates from severe cases of dermatophytosis in the United Kingdom, Ireland, France, Canada and India for the period 2018-2023, including the type strain from Japan. We used whole-genome sequencing to confirm 90 isolates were T. indotineae, and antifungal susceptibility testing indicated that over half of these (62%) were resistant to terbinafine (MIC ≥1 mg/L). Pairwise genetic distances showed very high identity with only 147 (1-414) SNPs separating isolates that were nested within a monophyletic phylogeny, supporting a single evolutionary origin of T. indotineae. That no clear geographic clustering of isolates was observed confirms the rapid transcontinental spread of T. indotineae from its likely centre of diversity in Asia. Genome-wide analyses identified multiple non-synonymous SNPs in SQLE (ERG1), the squalene epoxidase target of terbinafine, that were associated with terbinafine in vitro resistance ≥1 mg/L. However, five isolates exhibited high MIC values without SQLE mutations, suggesting the presence of alternative resistance mechanisms. Our findings highlight the importance of better genomic surveillance to understand and manage this severe and rapidly emerging terbinafine-resistant dermatophyte.

## Introduction

Dermatophytosis is the most common fungal infection globally affecting an estimated 20% – 25% of the world’s population (1–3). Infections are caused by dermatophytes, a ubiquitous and diverse group of keratinophilic filamentous fungi capable of invading human skin, hair and nails. Dermatophyte skin infections, also known as tinea or ringworm, are characterised by annular, pruritic lesions that easily spread, and are mainly attributed to Trichophyton rubrum and Trichophyton interdigitale (4). Secondary infection from other affected body sites or fomites can occur (5). Most cases of superficial dermatophytosis remain localized and respond well to standard topical or oral antifungal therapy, although onychomycosis (nail infection) caused by dermatophytes is generally classed as difficult to treat.

In the past decade, Trichophyton indotineae (previously Trichophyton mentagrophytes genotype VIII), a novel and frequently terbinafine-resistant dermatophyte species, has caused an epidemic of difficult-to-treat disease in the Indian subcontinent (6, 7). It is now a common infection, and often presents as widespread, inflammatory pruritic plaques on body, groin, thighs, pubic/genital area, buttocks, and the face (8, 9), usually manifesting as chronic, recurrent, or recalcitrant disease. Skin lesions often affect several body sites, and in some cases the possibility of sexual transmission was proposed (10). Most isolates of T. indotineae are clinically resistant to the topical allylamine drug terbinafine, the widely used standard first-line therapy, and frequently exhibit elevated in vitro MICs (MIC ≥1 mg/L) with a prevalence up to 75% according to certain studies (11). Infections thus often require subsequent prolonged treatment courses with itraconazole (12), although azole antifungal resistance can also occur. In vitro terbinafine resistance is associated with a phenylalanine-to-leucine amino acid change at codon 397 (F397L) along with other known alleles in the gene encoding the target of terbinafine, squalene epoxidase (ERG1/SQLE). It has been hypothesised that overuse of topical creams containing combinations of antifungals, antibacterials, and potent corticosteroids is the main driver leading to the evolution of terbinafine-resistant T. indotineae on the Indian subcontinent (13).

Since its discovery, infections caused by T. indotineae are increasingly reported worldwide (8, 11, 14, 15), and spread outside of the original endemic areas has been linked to international travel (16). However, there is now growing evidence of autochthonous acquisition of infection outside the previously established endemic regions (17, 18). Given the severity and drug-resistant phenotype associated with T. indotineae alongside a lack of understanding of the extent, ease and mode of transmission, there is an urgent need to better understand this species. Here we report results of whole-genome sequence (WGS) analysis of a panel of T. indotineae isolates sourced from six countries and three continents – Asia, Europe and North America – alongside their antifungal susceptibility patterns.

## Material and Methods

### Fungal isolates

103 Trichophyton human isolates during 2018-2023 were collected from United Kingdom (n = 25; 6 centres), France (n = 20; 1 centre), Canada (n = 19; 1 centre), Ireland (n = 1; 1 centre), and India (n = 31; 1 centre) (Supplementary Table 1). In addition, T. indotineae type-strain CBS 146623 isolated in 2019 in Japan (19) was included and sequenced. Isolates from India (A. Chowdhary) were subjected to DNA extraction and Illumina short-read whole genome sequencing prior to the start of this project and the reads were combined with the isolates sequenced in the current study. Case patient information for a subset of patients in France and Canada was previously described elsewhere (14, 15).

### Antifungal susceptibility testing (AFST)

For isolates collected from UK, Ireland and Canada, terbinafine antifungal susceptibility testing was determined according to the CLSI M38-A2 broth microdilution method (20). Minimum inhibitory concentration (MIC) for terbinafine was read at 80% inhibition of growth compared to the drug-free growth control after at least 96 hours of incubation. For isolates collected from France, the EUCAST E.Def 11.0 standard method (21) was used for AFST. Isolates with terbinafine MIC ≥1 mg/L were considered as non-wild-type (WT) and resistant, based on the distribution of MIC values of isolates within this study (Supplementary Figure 6). For Indian isolates AFST was carried out using the Clinical and Laboratory Standards Institute broth microdilution method (CLSI-BMD), using the M38 3^rd^ edition guidelines (22). The isolates were grown for 7 days on PDA at 27⁰C and conidial inoculum was prepared in saline containing Tween 80 by gently scraping the surface of mature colonies with a sterile cotton swab moistened with sterile physiological saline and allowed to settle. The final inoculum concentration was adjusted to twice the concentration needed for testing i.e., 1 × 10^3^ to 3 × 10^3^ CFU/ml counted by haemocytometer (CLSI). Antifungal drugs tested included allylamine viz. terbinafine (TRB; Synergene India, Hyderabad, India,), orotomide viz. olorofim, triazoles, viz. itraconazole (ITC; Lee Pharma, Hyderabad, India), voriconazole (VRC; Pfizer Central Research, Sandwich, Kent, United Kingdom), fluconazole (FLU; Sigma-Aldrich, Germany), posaconazole (POS; Sigma-Aldrich, Germany), isavuconazole (ISA; Sigma-Aldrich, Germany); imidazoles viz., luliconazole (LUZ, Emcure, Pune, India), and ketoconazole (KTC; Sigma-Aldrich); amphotericin B (AMB; Sigma-Aldrich) and griseofulvin (GRE; Sigma-Aldrich). Drug-free and mould-free controls were included, and microtiter plates were incubated at 30°C. Minimum inhibitory concentration (MIC) endpoints for all the drugs were defined as the lowest concentration that produced 80% inhibition of growth as read visually at 96h.

### DNA extraction and purification

Genomic DNA (gDNA) for isolates from Canada, UK and France were extracted as previously described (23). Briefly, fungal isolates were subcultured on Sabouraud glucose agar (SGA) plates supplemented with chloramphenicol and incubated at 28°C for 7-10 days until sporulation. Stock conidial suspensions were prepared by washing the surface of the SGA plates with 10 ml of sterile water containing 0.05% Tween 20. The conidial suspensions were filtered using Miracloth (EMD Chemicals, San Diego, CA, United States) to remove fungal hyphae, transferred to 50-ml sterile conical tubes, and centrifuged at maximum speed (10,000 × g) for 10 min. The supernatants were discarded, and the pellets were resuspended in 5 ml of sterile distilled water. The concentrations of the suspended conidial stocks were determined by counting the conidia using a haemocytometer chamber at ×400 magnification. Harvested conidia at concentrations of 2 × 10^8^/ml were subjected to DNA extraction. High molecular weight DNA was extracted with an optimized MasterPure™ Complete DNA and RNA purification kit (Lucigen, US) with an additional bead-beating step included. Harvested conidia were homogenized using 1.0 mm-diameter zirconia/silica beads (BioSpec Products, Bartlesville, OK) in a FastPrep-24 system (MP Biomedicals, Solon, OH) at 4.5 m/s for 45 s. Following a purification and concentration step using DNeasy Blood and Tissue kit (Qiagen, Germany), gDNA was quantified using a Qubit 2.0 fluorometer and dsDNA BR (double-stranded DNA, broad-range) assay kit (Life Technologies, Carlsbad, CA). Quality control of extracted gDNA samples prior to library preparation was performed using the TapeStation 2200 system (Agilent, Santa Clara, CA) and gDNA ScreenTape assays. gDNA libraries were constructed, normalised and indexed at Earlham Institute (UK) and run on a NovaSeq 6000 SP v1.5 flow cell to generate 150 bp paired-end reads. For Indian isolates, DNA was extracted using a column-based method with a QIAamp DNA minikit (Qiagen, Hilden, Germany) and was quantified by QUBIT 3 Fluorometer using dS DNA HS Dye. WGS libraries were prepared using NEBNext ultra II DNA FS kit (New England Biolabs, Ipswich, MA, USA). In brief, 200 ng of intact DNA was enzymatically fragmented by targeting 200–300 bp fragments sizes followed by purification using AMPure beads (Beckman Coulter Life Sciences, Indianapolis, IN, USA). For sequencing, the libraries were normalized to 10 nMol/L concentration and pooled together at equal volumes. Further, the library pools were denatured using freshly prepared 0.2 N NaOH for cluster generation on cBOT and sequenced on Illumina HiSeq 4000.

### Whole genome sequencing and analysis, including single-nucleotide polymorphism identification

Sequence alignment and quality control. Raw Illumina reads were mapped to the T. indotineae reference genome (GenBank GCA_023065905.1; strain TIMM20114) using the Burrows Wheeler Aligner (BWA) MEM algorithm v0.7.17 (24). Reference genomes used for the alignment of T. interdigitale and T. mentagrophytes were GCA_019359935.1 and GCA_003664465.1, respectively. The output SAM files were then converted to compressed BAM format, which underwent sorting and indexing using SAMTools v1.6.1 (25). Picard v2.27.4 was used to identify and mark duplicated reads (26). GATK HaplotypeCaller version 4.2 was used to call SNPs, insertions, and deletions, excluding repetitive regions as identified above (27, 28). Quality control measures were implemented to filter out low-confidence SNPs if they met at least one of the following criteria: “QualityByDepth < 2.0 || FisherStrand > 60.0 || MappingQuality < 40.0 || MQRankSum < -12.5 || ReadPosRankSum < -8.0 || StrandOddsRatio > 4.0”. The T. indotineae reference genome was annotated by identifying repeat content using Repeatmodeler v2.0.3 (29) (engine NCBI) with CD-HIT (30), ReCon v1.08 (31), RMBlast v2.11.0, Tandem Repeats Finder v4.09 (32), RepeatScout v1.06 (33) and RepeatMasker v4.1.2-p1 (34). The output of Repeatmodeller was then used as a library for RepeatMasker to identify all instances of repeats and soft mask the genome (parameter - small). Gene annotation on the repeat masked assembly was achieved using the Braker2 (35) pipeline (parameters –fungus –softmasking), which uses Samtools v0.1.19-44428cd (25), Bamtools v2.4.0 (36), Diamond v2.0.4 (37), Genemark-ET v4.15 (38), and Augustus v3.2.1 (39). The pipeline identified 5,451 protein-coding genes, and BUSCO assessment resulted in 90.8% complete BUSCOs. The full genome annotation for the T. indotineae reference genome is available at 10.6084/m9.figshare.27275004. SnpEff version 5.1 was used to map SNPs to genes and infer functional annotation (40). The reference T. indotineae genome and the annotation file in gtf format with the specified options (-gtf22, -noGenome, -noCheckCds, -noCheckProtein) were used to construct a custom database for annotation. Annotated sequences were interrogated for non-synonymous SNPs (nsSNPs) in the SQLE coding area and ERG11 gene that could be linked to terbinafine resistance. The 1 kb upstream nucleotide sequence of SQLE was assessed for the presence of promoter tandem repeats using TRF v4.07b (32).

### Population and mating type analysis

Genetic similarity of isolates was investigated based on SNP data using principal component analysis (PCA) using ‘dudi.pca’ from the ‘ade4’ v1.7-22 package in R v4.3.2. Calculation of Weir & Cockerham (41) F_ST_values for between-country diversity were conducted in R v4.4.1 using ‘hierfstat’ v0.5-11 (42). Mating type was reported for all T. indotineae isolates based on presence of the HMG gene (GenBank: MW551758.1) and/or α-box mating type locus (GenBank: OQ536564.1) sequences. The presence of the α-box indicates the MAT1-1 mating type, whereas the presence of the HMG gene indicates the MAT1-2 mating type.

### Phylogenetic analysis

MultiFASTA files containing whole genome SNP calls were used to construct maximum-likelihood phylogenetic trees with Randomized Axelerated Maximum Likelihood (RAxML-HPC) version 8.2.12 (43), using the GTRCAT approximation of rate heterogeneity and 500 bootstrap replicates. Resulting phylogenies were visualised using ggtree version 3.1.14 (44) in R version 4.3.2. The best-fitting RAxML phylogeny and associated isolate metadata can be accessed as a Microreact project (45) at: https://microreact.org/project/wYSoQggWejRkhX2LnsMtWt-trichophyton (Supplementary Figure S1).

### Identifying loci associated with terbinafine resistance

In order to assess which loci were associated with terbinafine resistance, we performed genome-wide association (GWA) tests using treeWAS v1.0 (46). A nucleotide alignment and phylogeny of all confirmed T. indotineae, T. mentagrophytes, and T. interdigitale isolates was used, generated as described above, to achieve sufficient statistical power. MIC information was available for all isolates, which were defined as either susceptible (MIC <1 mg/L) or resistant (MIC ≥1 mg/L). treeWAS was performed in R v4.3.2 with a p-value cutoff of 0.05 for three tests of association (subsequent, terminal, and simultaneous) between terbinafine resistant and susceptible phenotypes as a discrete vector.

### Data availability

All raw genomic data are available under the Project Accession PRJEB75499. Scripts are available at https://github.com/mycologenomics/Trichophyton

## Results

### Relationship of global T. indotineae isolates using WGS

A collection of clinical Trichophyton isolates sourced from six countries and three continents were whole genome sequenced to assess the genetic basis of terbinafine (TERB) resistance and population structure of T. indotineae in relation to the rest of the Trichophyton genus. Isolates had been recovered from patients with provisional diagnosis of Trichophyton infection and provisionally identified initially either by ITS sequencing and/or phenotypic characterisation. The collection encompassed 103 isolates from various geographical locations, including the type strain from Japan (Figure 1a, Supplementary Table 1).

**Figure 1:**
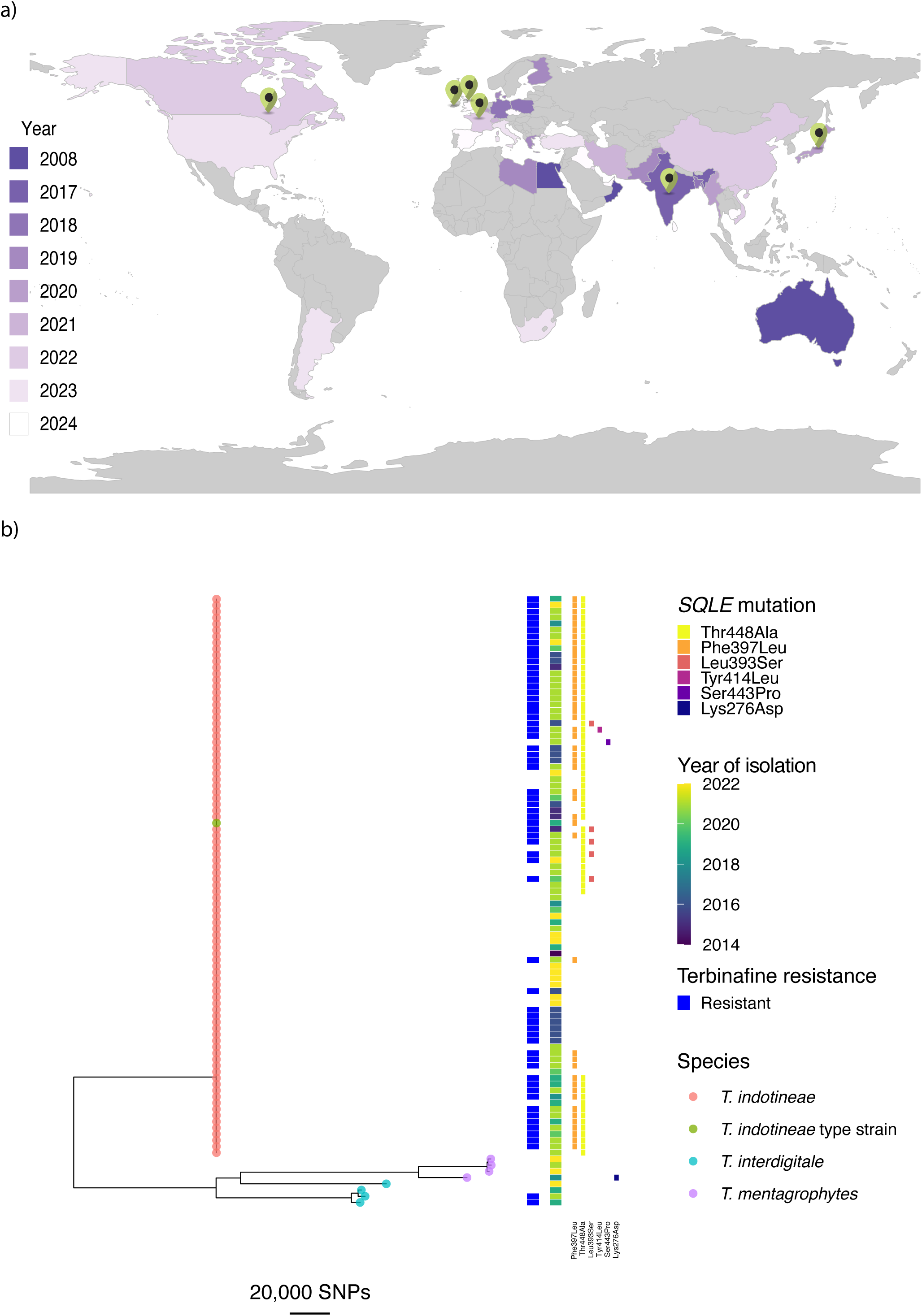
a) Confirmed global reports of T. indotineae indicated by year of report. Green points indicate location of sampling for isolates included in this study b) Genetic relationships among 102 isolates inferred using a maximum likelihood phylogeny for isolates identified as T. mentagrophytes, T. interdigitale and T. indotineae, including the T. indotineae type strain. Isolates with raised terbinafine MIC values, and therefore considered resistant, are indicted. Presence of mutations conferring amino acid substitutions in SQLE are also indicated. Branch lengths represent the average number of SNPs along that branch.

The median percentage of reads mapped to reference was 97.54%, (range: 85.33% to 99.09%). The sequences from most of the isolates (96/103) could be successfully mapped to the T. indotineae reference genome (successful mapping threshold exceeding 90%), indicating reliable data for downstream analyses. Four isolates displayed low percentage mapping to the T. indotineae reference genome (Tri_36, 93, 95 and 97, range: 3.90-6.24%). These four isolates also displayed low percentage mapping to the T. interdigitale (5.70-6.66%) and T. mentagrophytes (range: 5.94-7.30%) reference genomes (Supplementary Table 2). The raw sequence data was queried using BLASTn v2.15.0; Tri_36, 93 and 95 were identified as Cordyceps fumosorosea, and Tri_97 was identified as Emydomyces testavorans. Interestingly, the Tri_97 isolate also displayed raised TERB MIC (4 mg/L). These isolates were excluded from further analysis, due to previous misidentification.

The number of high-confidence SNPs called ranged from 20 SNPs to 130,617 SNPs, with a median of 115 SNPs. Isolates with more than 100,000 SNPs called showed higher percentage mapping to either T. interdigitale or T. mentagrophytes reference genomes, compared to mapping against T. indotineae (Supplementary Table 2). These isolates (n = 8; Tri_02, Tri_04, Tri_18, Tri_39, Tri_80, Tri_84, Tri_85, Tri_94) also displayed high bootstrap support (100%) and clustered with type genomes for T. interdigitale or T. mentagrophytes, and were therefore identfied as either T. interdigitale or T. mentagrophytes (Supplementary Figure 2). In total, 90 isolates mapped with high fidelity to the reference T. indotineae genome with low numbers of filtered SNP differences (range: 20-346), plus the type strain, confirming their correct identification as this species. Figure 1b illustrates the divergence of the eight non-T. indotineae isolates along an extended branch, and further forming two different clusters within the Trichophyton mentagrophytes/Trichophyton interdigitale species complex (TMTISC). Principal Component Analysis (PCA) confirmed separation of T. interdigitale and T. indotineae from T. mentagrophytes on PC2 (Supplementary Figure S3a), with isolates from all countries occupying the same space of genetic diversity (Supplementary Figure S3b); isolates from Ireland, India and Japan were almost indistinguishable.

On average (mean), 147 SNPs separate any pair of T. indotineae isolates (median: 121 SNPs; range: 1-414 SNPs). In contrast, any T. indotineae isolate is, on average, separated from T. interdigitale or T. mentagrophytes isolates by 231,767 and 290,500 SNPs, respectively, indicating significant divergence of the TMTISC complex. We included the T. indotineae type strain CBS 146623 in this analysis, and isolates identified as T. indotineae were on average separated by 102 SNPs from this type strain (Figure 1b). All T. indotineae isolates contained the HMG gene, indicating the presence of MAT1-2 mating type only, confirming earlier studies (47).

We observed no clustering of isolates by sampling location or time (Figure 1b); a PCA based on whole genome SNPs for only the confirmed T. indotineae isolates showed separation of Irish and Canadian isolates based on PC2 (Figure 3b). However, isolates from all countries could be seen to be drawn from the same population. Two lineages containing mainly terbinafine-resistant isolates sampled from UK, Canada, and France were observed (Figure 3a); these branches were highly supported (range: 97-100%; Supplementary Figure S4), indicating potentially unsampled diversity in Asia leading to the emergence of these lineages.

Within-country diversity was assessed by average (mean) pairwise SNP distances where more than one isolate was sampled (therefore, excluding Japan and Ireland; Supplementary Figure S5a; Supplementary Table S3). Generally, average pairwise SNP distances were similar regardless of the country of origin (Supplementary Table S3); we observed clusters of closely related isolates sampled from Canada, France and UK, whereas larger genetic diversity was observed in India (Supplementary Figure 5a). Comparison of common and unique SNPs amongst isolates within Canada, UK, France and India revealed less than 10% of SNPs were common to all four countries (Supplementary Figure S5b); yet each country had, on average, 262 unique SNPs (range: 200 – 375). Between country diversity was assessed via the fixation index F_ST_ (Supplementary Figure S5c; Supplementary Table S4); values were approaching zero in all comparisons, indicating complete sharing of genetic material. However, F_ST_for India and Canada, and India and the UK, were higher (0.038 and 0.024, respectively), indicating some isolation between T. indotineae sampled in these countries.

### Antifungal drug resistance

AFST was performed for all T. indotineae and non-indotineae isolates in this study. Using an MIC breakpoint of ≥1 mg/L, 54 T. indotineae isolates (53%) were therefore determined to be resistant to terbinafine, with 20 isolates (20%) displaying an MIC of 16 or higher (Supplementary Figure S6). We therefore sought to identify the genetic basis of terbinafine resistance by performing genome-wide association using treeWAS, which accounts for confounding effects of clonality and genetic structure. treeWAS identified a single significant locus (p-value = 2.225^e-308^) that mapped to the SQLE open reading frame, resulting in the missense mutation causing a change from phenylalanine to leucine at position 397 (Phe397Leu). This statistically significant locus was present in 34 of the resistant isolates, which displayed terbinafine MIC ranges from 1 to ≥16 mg/L. We therefore sought to further characterise the known mutations in SQLE, which encodes the target of terbinafine, and ERG11, encoding an enzyme targeted by other antifungal drug classes.

We examined the extent that polymorphisms within SQLE were associated with resistance to terbinafine among the 90 T. indotineae isolates by identifying non-synonymous SNPs (nsSNPs) conferring amino acid changes. Among the T. indotineae isolates, six nsSNPs were identified in SQLE that conferred amino acid substitutions: Leu393Ser, Leu393Phe, Phe397Leu, Tyr414His, Ser443Pro, and Thr448Ala. Notably, Thr448Ala (n = 60) was the most prevalent substitution and 80% of isolates with this change exhibited raised MIC to terbinafine (Figure 2). However, 19 isolates contained the Thr448Ala mutation alone, with 63.16% being non-resistant and 36.84% being resistant. This finding suggests that the Thr448Ala substitution is not sufficient to confer terbinafine resistance alone. The second most common substitution was Phe397Leu (n = 45), and 100% of isolates with this allele demonstrated MIC values of ≥1 mg/L and therefore resistance to terbinafine (Figure 2). The substitution Ser443Pro (n = 1), Leu393Phe (n = 1), and Leu393Ser (n = 4) were all found in combination with other substitutions, notably Thr448Ala, while Tyr414His (n = 1) was found in conjunction with both Thr448Ala and Phe397Leu (Figure 3a).

**Figure 2:**
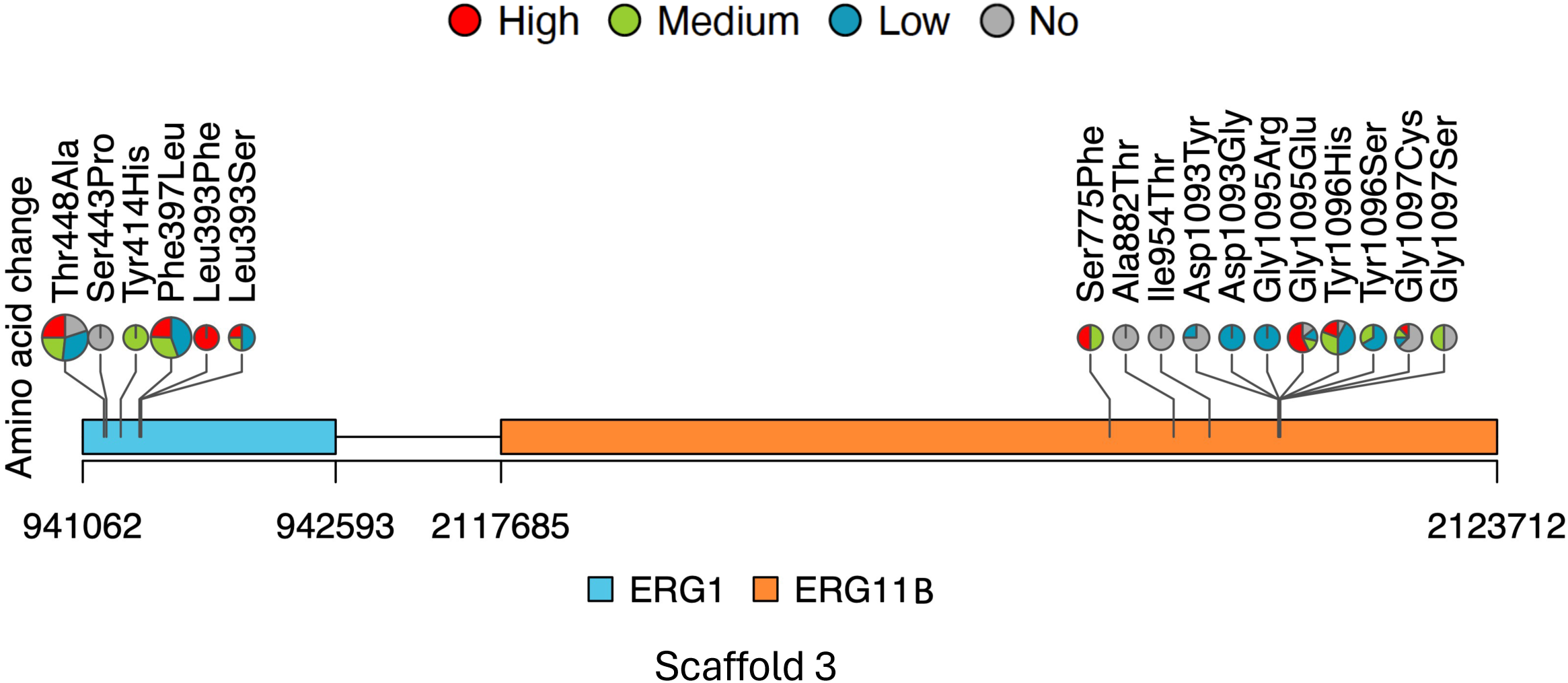
Amino acid location changes and corresponding SNP changes in SQLE and ERG11 genes, present in the third scaffold (JAJVHL0100000003.1) of the ASM2306590v1 assembly. Pie chart size represents the frequency of the SNP occurrence.

**Figure 3:**
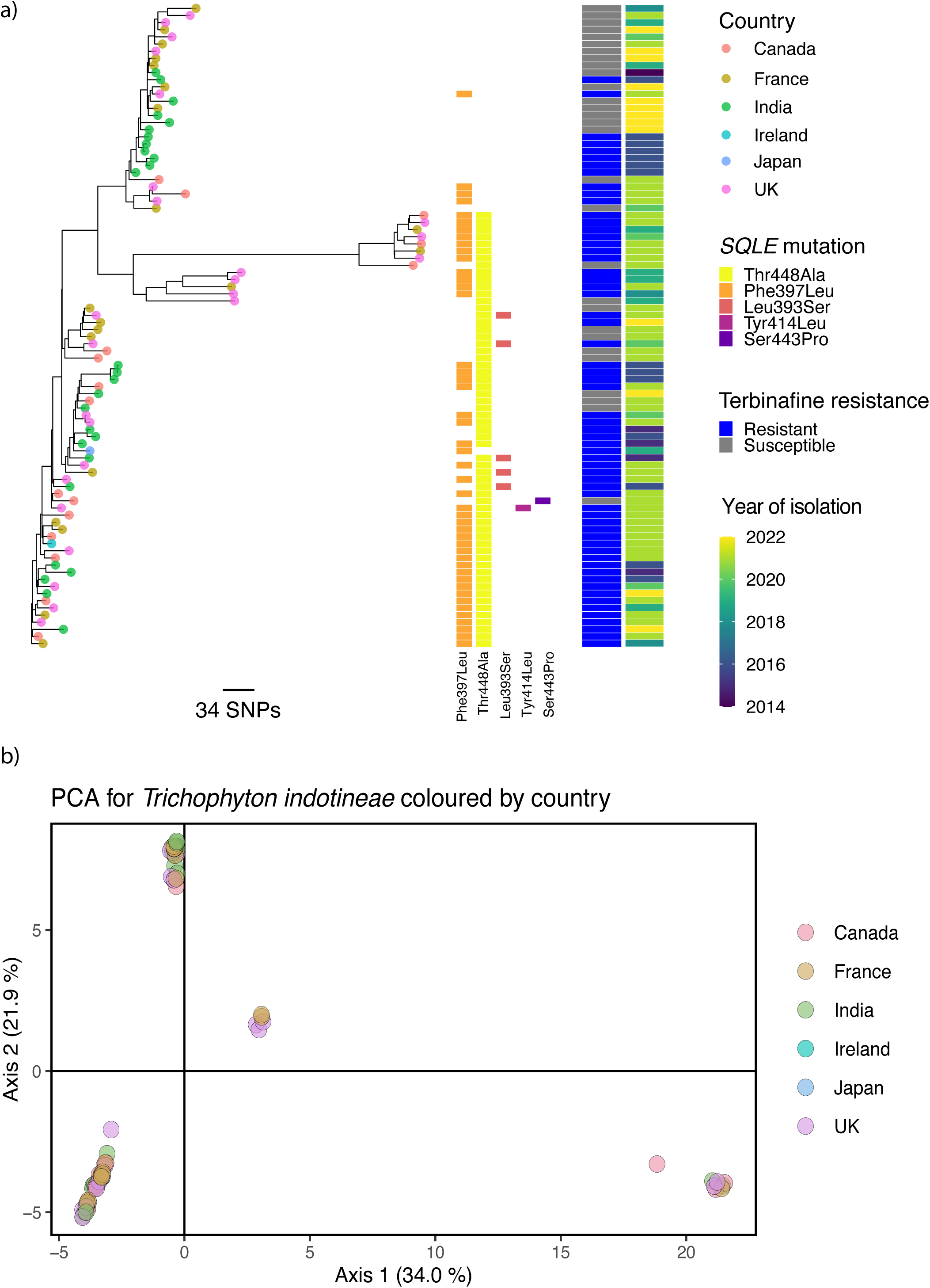
a) Maximum-likelihood phylogeny of all confirmed T. indotineae isolates. Tip points indicate country of origin. Isolates with raised terbinafine MICs, and therefore resistant, are shown in blue. SQLE amino acid substitutions are also shown b) Principal components analysis (PCA) for all confirmed T. indotineae isolates shows some separation of Irish and Canadian isolates based on PC2. However, isolates from all countries could be seen to be drawn from the same population.

Although certain allelic combinations were limited in occurrence, it is noteworthy that majority of isolates with the Tyr414His, Leu393Phe, and Leu393Ser combination of substitutions exhibited resistance to terbinafine (Figure 3a). Whilst the Leu393Phe (L398F) substitution has been confirmed to be involved in terbinafine resistance in T. rubrum (48), further investigation is required to understand the significance of these genotypes in contributing to terbinafine resistance. The Thr448Ala allele was of further interest as it appeared in both T. indotineae and non-T. indotineae isolates. This intriguing discovery raises further questions about the potential role of the Thr448Ala mutation in relation to terbinafine resistance.

Finally, almost a third of T. indotineae isolates (n = 28; 30.43%) had no SNP identified in SQLE. More than 70% of these SQLE WT isolates (20/28; 71.43%) were susceptible to terbinafine. Eight out of these 28 WT isolates exhibited high levels of in vitro resistance to terbinafine with MICs >16 mg/L, indicating the presence of alternative resistance mechanisms that do not rely on changes in SQLE. In the exploration of additional resistance mechanisms, an examination of the promoter region of the SQLE gene revealed no tandem repeats.

As ERG11 is the primary target of azole antifungal drugs, we aimed to characterise mutations within this gene, and to observe any associations with mutations within ERG1. T. indotineae is thought to contain two paralogs of ERG11, ERG11A and ERG11B (49). A single nsSNP conferring a Arg817Cys mutation in ERG11A was present in five isolates (Tri-15, 16, 44, 69, and 73), none of which exhibited any terbinafine resistance. Of the remaining T. indotineae isolates, all exhibited a wild-type ERG11A sequence, suggesting that ERG11A is not involved in terbinafine resistance. Next, we analysed SNPs in the ERG11B sequence of T. indotineae, identifying Ser775Phe, Ala882Thr, Ile954Thr, Asp1093Tyr, Asp1093Gly, Gly1095Arg, Gly1095Glu, Tyr1096His, Tyr1096Ser, Gly1097Cys, and Gly1097Ser (Figure 2). Notably, 21.67% (13/60) of isolates with the Thr448Ala in ERG1 also possessed the Gly1095Glu mutation in ERG11B. Similarly, 43.33% (26/60) with Thr448Ala in ERG1 also contained the Tyr1096His mutation in ERG11B. These overlapping mutations suggest potential interactions between ERG1 and ERG11B, possibly contributing to the development of terbinafine resistance. Indeed, statistical analyses highlighted significant associations between mutations in SQLE (p-value = 0.00469; Chi-squared test) and ERG11B (p = 1.08^e-04^) and in vitro terbinafine resistance; however, we did not systematically assess these isolates for sensitivity to any azole antifungal drug. This observation underscores the need for further investigation to fully elucidate the complex genetic factors driving terbinafine resistance in individual isolates in T. indotineae.

## Discussion

T. indotineae has recently become a significant public health concern in a number of countries by causing difficult to treat disease, and displaying high levels of terbinafine antifungal drug resistance (7). Here, we report isolates from clinical Trichophyton infections from Canada, Ireland, UK, France, and India. The whole genome sequencing of 103 clinical Trichophyton isolates, including the type strain from Japan, provided significant insights into the genetic basis of antifungal drug resistance and the population structure of T. indotineae. Our findings highlight both the genetic diversity and the underlying mechanisms potentially contributing to terbinafine resistance, underscoring the complex nature of antifungal drug resistance in this emerging fungal pathogen.

Our WGS data reveals a high degree of genetic fidelity among 90 T. indotineae isolates, as demonstrated by the high percentage of reads mapped to the reference genome and low numbers of SNPs. The divergence observed in eight non-T. indotineae isolates emphasises the importance of accurate species identification in understanding the epidemiology and resistance patterns of fungal pathogens.

The genetic diversity of T. indotineae was characterised by an average of 147 SNPs separating any pair of isolates, with no significant clustering by geographic location or time. This lack of clustering suggests a widespread and genetically homogenous distribution of T. indotineae across different regions, consistent with previous findings that indicate a high level of genetic similarity among isolates from disparate locations (17). However, assessment of within-country and between-country diversity using pairwise SNPs and F_ST_ revealed clusters of closely related isolates within Canada, France and UK but not India. This suggests that this study under-sampled one likely source of T. indotineae (India), and has therefore not observed the true extent of genetic diversity. Future efforts to undertake large-scale global genomic surveillance would resolve this issue.

Antifungal susceptibility testing revealed that over half of the T. indotineae isolates in this study exhibited in vitro resistance to terbinafine, with a high proportion showing high MIC values (>16 mg/L). The genome-wide association analysis identified a single significant locus associated with terbinafine resistance, yet this could not fully explain the resistance observed, pointing to the involvement of additional genetic factors. Multiple nsSNPs were identified in SQLE, particularly the prevalent Thr448Ala mutation, and the previously reported Phe397Leu mutations (17, 50, 51). However, the presence of either of these mutations alone did not uniformly confer resistance, suggesting other mutations or genetic factors may modulate its impact. Interestingly, a subset of T. indotineae isolates exhibited high levels of terbinafine resistance despite the absence of SNPs in SQLE, indicating the existence of alternative resistance mechanisms. Whilst our findings are consistent with other studies that also found T. indotineae isolates were wildtype for ERG11A, we did not find previously identified ERG11B mutations (49). Our analysis also suggests an association between mutations in SQLE and ERG11B and terbinafine resistance, pointing to a potential synergistic interaction between these genes in conferring resistance.

Highly supported lineages identified by sampling isolates within the UK, Canada and France suggest under-sampling of the full extent of genetic diversity in Asia, despite the majority of isolates in this study originating in India. Currently, under-sampled areas include Bangladesh and Pakistan. Ultimately, our findings highlight the necessity for ongoing genomic surveillance and detailed genetic analyses to monitor the spread and evolution of antifungal drug resistance in T. indotineae. Future research should focus on elucidating the full spectrum of genetic changes contributing to terbinafine resistance, including the potential roles of regulatory elements and non-coding regions. Linking this work with a newly established international database for drug-resistant tinea, established by the International League of Dermatological Societies (ILDS) in collaboration with the American Academy of Dermatology (AAD) would help to further our understanding of this growing problem.

## Author contributions

Isolate collection and preparation: RS, SD, JAS, AC, RCB, JL, SS, RH, AB, SH, RL, RG

Sequencing: JAS, AC, MCF, JR, AK

Data analysis: JR, STH, RF

Manuscript preparation: JR, STH, AA, MCF

## Funding sources

RAF is supported by a Wellcome Trust Career Development Award (225303/Z/22/Z). JR is supported by a Wellcome Trust ISSF Springboard Fellowship.

MCF is supported by the Wellcome Trust (219551/Z/19/Z) and is a fellow of the Canadian Institute for Advanced Research (CIFAR).

AC is support be Science and Engineering Research Board (SERB File No. CRG/2020/001735) Department of Science and Technology, Government of India, and is a fellow of the Canadian Institute for Advanced Research (CIFAR).

