## Supplementary Information for "Genomic epidemiology of emerging terbinafine-resistant Trichophyton indotineae"

**Supplementary Table S1: *Trichophyton* isolate metadata, including terbinafine MIC (mg/L) and associated *SQLE* (*ERG1*) SNPs.** NA = data not available. Isolates highlighted in grey signify non-*indotineae* *Trichophyton* isolates. MIC values considered resistant (MIC ≥1mg/L) are given in bold typeface.

| Study number | Country | Specimen type/site | Year collected | Identification | Travel history | TERB MIC (mg/L) | *SQLE* SNPs | Culture collection ID (where available) |
| --- | --- | --- | --- | --- | --- | --- | --- | --- |
| Tri-01 | UK | Skin, buttock | 2018 | *T. indotineae* |  | **1** | Phe397Leu, Thr448Ala |  |
| Tri-02 | UK | Skin, unknown | 2019 | *T. interdigitale* |  | 0.125 |  |  |
| Tri-03 | UK | Skin, groin | 2019 | *T. indotineae* |  | **4** | Phe397Leu, Thr448Ala |  |
| Tri-04 | UK | Skin, arm | 2019 | *T. interdigitale* |  | **8** |  |  |
| Tri-05 | UK | Skin, back | 2019 | *T. indotineae* | India | **>16** | Phe397Leu, Thr448Ala |  |
| Tri-06 | UK | Skin, torso | 2019 | *T. indotineae* |  | **4** | Phe397Leu, Thr448Ala |  |
| Tri-07 | UK | Skin, arm | 2020 | *T. indotineae* |  | **8** | Phe397Leu, Thr448Ala |  |
| Tri-08 | UK | Skin, Leg/foot | 2020 | *T. indotineae* |  | **2** | Phe397Leu, Thr448Ala |  |
| Tri-09 | UK | Skin, buttock | 2020 | *T. indotineae* |  | **2** | Phe397Leu, Thr448Ala |  |
| Tri-10 | UK | Skin, groin | 2021 | *T. indotineae* |  | **2** | Phe397Leu, Thr448Ala |  |
| Tri-11 | UK | Skin, buttock | 2021 | *T. indotineae* |  | **2** | Phe397Leu, Thr448Ala |  |
| Tri-12 | UK | Skin, axilla | 2021 | *T. indotineae* |  | **4** | Phe397Leu, Tyr414Leu, Thr448Ala |  |
| Tri-13 | UK | Skin, thigh | 2021 | *T. indotineae* |  | **8** | Phe397Leu |  |
| Tri-14 | UK | Skin, buttock | 2019 | *T. indotineae* |  | 0.125 | Thr448Ala |  |
| Tri-15 | UK | Skin, groin | 2019 | *T. indotineae* |  | <0.03 | Wildtype |  |
| Tri-16 | UK | Skin, groin | 2020 | *T. indotineae* |  | 0.06 | Wildtype |  |
| Tri-17 | UK | Skin, groin | 2020 | *T. indotineae* |  | **2** | Leu393Ser, Thr448Ala |  |
| Tri-18 | UK | Skin, face | 2021 | *T. mentagrophytes* |  | <0.03 |  |  |
| Tri-19 | UK | Skin, back | 2021 | *T. indotineae* | India | **2** | Phe397Leu |  |
| Tri-20 | UK | Skin, unknown | 2021 | *T. indotineae* | India | **2** | Phe397Leu, Thr448Ala |  |
| Tri-21 | Canada | Skin, buttock | 2021 | *T. indotineae* |  | **2** | Phe397Leu, Thr448Ala | UAMH12246 |
| Tri-22 | Canada | Skin, buttock | 2021 | *T. indotineae* |  | **2** | Phe397Leu, Thr448Ala | UAMH12247 |
| Tri-23 | Canada | Skin, unknown | 2021 | *T. indotineae* |  | **2** | Phe397Leu, Thr448Ala | UAMH12248 |
| Tri-24 | Canada | Skin, hands | 2021 | *T. indotineae* |  | <0.03 | Thr448Ala | UAMH12249 |
| Tri-25 | Canada | Skin, buttock | 2021 | *T. indotineae* |  | **2** | Phe397Leu, Thr448Ala | UAMH12250 |
| Tri-26 | Canada | Skin, thigh | 2021 | *T. indotineae* |  | 0.06 | Ser443Pro, Thr448Ala | UAMH12251 |
| Tri-27 | Canada | Skin, back | 2021 | *T. indotineae* |  | **2** | Phe397Leu, Thr448Ala | UAMH12335 |
| Tri-28 | Canada | Skin, back | 2021 | *T. indotineae* |  | <0.03 | Wildtype | UAMH12336 |
| Tri-29 | Canada | Skin, leg | 2021 | *T. indotineae* |  | <0.03 | Thr448Ala | UAMH12337 |
| Tri-30 | Canada | Skin, neck | 2021 | *T. indotineae* |  | **2** | Phe397Leu | UAMH12338 |
| Tri-31 | Canada | Skin, abdomen | 2021 | *T. indotineae* |  | **2** | Phe397Leu, Thr448Ala | UAMH12339 |
| Tri-32 | Canada | Skin, unknown | 2021 | *T. indotineae* |  | **2** | Phe397Leu, Thr448Ala | UAMH12340 |
| Tri-33 | Canada | Skin, thigh | 2021 | *T. indotineae* |  | **1** | Phe397Leu, Thr448Ala | UAMH12341 |
| Tri-34 | Canada | Skin, buttock | 2021 | *T. indotineae* |  | 0.06 | Thr448Ala | UAMH12342 |
| Tri-35 | Canada | Skin, pubic area | 2021 | *T. indotineae* |  | **2** | Phe397Leu, Thr448Ala | UAMH12343 |
| Tri-36 | Canada | Skin, unknown | 2021 | Non*-Trichophyton* |  | <0.03 |  | UAMH12344 |
| Tri-37 | Canada | Skin, buttock | 2021 | *T. indotineae* |  | 0.06 | Thr448Ala | UAMH12345 |
| Tri-39 | UK | Nail | 2021 | *T. interdigitale* |  | **1** |  |  |
| Tri-40 | UK | Skin, unknown | 2021 | *T. indotineae* |  | **2** | Phe397Leu |  |
| Tri-41 | UK | Skin, unknown | 2021 | *T. indotineae* |  | **16** | Phe397Leu, Thr448Ala |  |
| Tri-42 | UK | Skin, arm | 2021 | *T. indotineae* |  | **8** | Phe397Leu, Thr448Ala |  |
| Tri-43 | UK | Skin, groin | 2021 | *T. indotineae* |  | **4** | Phe397Leu, Thr448Ala |  |
| Tri-44 | UK | Skin, unknown | 2021 | *T. indotineae* |  | 0.125 | Wildtype |  |
| Tri-46 | UK | Skin, unknown | 2021 | *T. indotineae* |  | **4** | Phe397Leu, Thr448Ala |  |
| Tri-47 | Ireland | Skin, leg | 2021 | *T. indotineae* |  | **4** | Phe397Leu, Thr448Ala |  |
| Tri-48 | UK | Skin, torso | 2021 | *T. indotineae* |  | **1** | Leu393Ser, Thr448Ala |  |
| Tri-49 | UK | Skin, buttock | 2022 | *T. indotineae* | India | <0.03 | Wildtype |  |
| Tri-52 | France | Skin, groin | 2021 | *T. indotineae* | Bangladesh | **4** | Phe397Leu, Thr448Ala |  |
| Tri-53 | France | Skin, groin | 2018 | *T. indotineae* | India | **4** | Phe397Leu, Thr448Ala |  |
| Tri-54 | France | Skin, thigh | 2019 | *T. indotineae* | Bangladesh | **4** | Phe397Leu, Thr448Ala |  |
| Tri-56 | France | Skin, arm | 2018 | *T. indotineae* | India | <0.03 | Wildtype |  |
| Tri-57 | France | Skin, groin | 2019 | *T. indotineae* | Sri Lanka | 0.03 | Wildtype |  |
| Tri-58 | France | Skin, arm | 2021 | *T. indotineae* | India | **16** | Phe397Leu, Thr448Ala |  |
| Tri-59 | France | Skin, groin | 2020 | *T. indotineae* | Bangladesh | 0.03 | Wildtype |  |
| Tri-61 | France | Skin, thigh | 2021 | *T. indotineae* | Thailand | **16** | Phe397Leu, Thr448Ala |  |
| Tri-63 | France | Skin, groin | 2021 | *T. indotineae* | Bangladesh | 0.125 | Thr448Ala |  |
| Tri-64 | France | Skin, thigh | 2021 | *T. indotineae* | India | **16** | Phe397Leu, Thr448Ala |  |
| Tri-66 | France | Skin, trunk | 2021 | *T. indotineae* | Bangladesh | **16** | Leu393Ser, Thr448Ala |  |
| Tri-67 | France | Skin, groin | 2021 | *T. indotineae* |  | 0.03 | Thr448Ala |  |
| Tri-68 | France | Skin, thigh | 2021 | *T. indotineae* | Bangladesh | 0.03 | Thr448Ala |  |
| Tri-69 | France | Skin, buttock | 2021 | *T. indotineae* | Bangladesh | 0.03 | Wildtype |  |
| Tri-70 | France | Skin, groin | 2021 | *T. indotineae* |  | **16** | Phe397Leu, Thr448Ala |  |
| Tri-71 | France | Skin, trunk | 2022 | *T. indotineae* | Afghanistan | 0.06 | Wildtype |  |
| Tri-73 | France | Skin, trunk | 2022 | *T. indotineae* | Bangladesh | 0.06 | Wildtype |  |
| Tri-74 | France | Skin, buttock | 2022 | *T. indotineae* | Bangladesh | 0.03 | Wildtype |  |
| Tri-77 | France | Skin, groin | 2022 | *T. indotineae* | Bangladesh | **2** | Thr448Ala |  |
| Tri-78 | France | Skin, axilla | 2022 | *T. indotineae* | Bangladesh | <0.03 | Wildtype |  |
| Tri-80 | France | Skin, face | 2018 | *T. mentagrophytes* |  | 0.03 | Lys276Asp |  |
| Tri-84 | France | Skin, buttock | 2022 | *T. mentagrophytes* | None | <0.03 |  |  |
| Tri-85 | France | Skin, pubic area | 2022 | *T. mentagrophytes* |  | <0.03 |  |  |
| Tri-93 | India | Skin, arm | 2022 | Non*-Trichophyton* |  | 0.25 |  |  |
| Tri-94 | India | Skin, face | 2022 | *T. interdigitale* |  | 0.125 |  |  |
| Tri-95 | India | Skin, abdomen | 2022 | Non*-Trichophyton* |  | 0.5 |  |  |
| Tri-96 | India | Skin, chest | 2022 | *T. indotineae* |  | **4** | Phe397Leu, Thr448Ala |  |
| Tri-97 | India | Skin, face | 2022 | Non*-Trichophyton* |  | **4** |  |  |
| Tri-98 | India | Skin, neck | 2022 | *T. indotineae* |  | 0.25 | Wildtype |  |
| Tri-99 | India | Nail | 2022 | *T. indotineae* |  | 0.03 | Wildtype |  |
| Tri-100 | India | Skin, abdomen | 2022 | *T. indotineae* |  | **8** | Phe397Leu, Thr448Ala |  |
| Tri-101 | India | Skin, face/trunk | 2022 | *T. indotineae* |  | 0.06 | Thr448Ala |  |
| Tri-102 | India | Skin, groin | 2022 | *T. indotineae* |  | 0.06 | Wildtype |  |
| Tri-103 | India | Skin, groin/trunk | 2021 | *T. indotineae* |  | 0.06 | Thr448Ala |  |
| Tri-104 | India | Skin, groin | 2022 | *T. indotineae* |  | 0.06 | Wildtype |  |
| Tri-105 | India | Skin, trunk | 2014 | *T. indotineae* |  | 0.5 | Wildtype |  |
| Tri-106 | India | Skin, groin | 2015 | *T. indotineae* |  | **1** | Phe397Leu, Thr448Ala |  |
| Tri-107 | India | Skin, groin | 2015 | *T. indotineae* |  | **>16** | Thr448Ala |  |
| Tri-108 | India | Skin, trunk | 2015 | *T. indotineae* |  | **>16** | Leu393Ser, Thr448Ala |  |
| Tri-109 | India | Skin, abdomen | 2015 | *T. indotineae* |  | **16** | Phe397Leu, Thr448Ala |  |
| Tri-110 | India | Skin, trunk | 2016 | *T. indotineae* |  | **>16** | Phe397Leu, Thr448Ala |  |
| Tri-111 | India | Skin, groin | 2016 | *T. indotineae* |  | **>16** | Wildtype |  |
| Tri-112 | India | Skin, groin | 2016 | *T. indotineae* |  | **>16** | Wildtype |  |
| Tri-113 | India | Skin, groin | 2016 | *T. indotineae* |  | **>16** | Wildtype |  |
| Tri-114 | India | Skin, unknown | 2016 | *T. indotineae* |  | **>16** | Phe397Leu, Thr448Ala |  |
| Tri-115 | India | Skin, abdomen | 2016 | *T. indotineae* |  | **>16** | Leu393Phe, Thr448Ala |  |
| Tri-116 | India | Skin, groin | 2016 | *T. indotineae* |  | **1** | Wildtype |  |
| Tri-117 | India | Skin, trunk/groin | 2016 | *T. indotineae* |  | **>16** | Wildtype |  |
| Tri-118 | India | Skin, trunk | 2016 | *T. indotineae* |  | **>16** | Wildtype |  |
| Tri-119 | India | Skin, groin | 2016 | *T. indotineae* |  | **1** | Phe397Leu, Thr448Ala |  |
| Tri-120 | India | Skin, groin | 2016 | *T. indotineae* |  | **>16** | Phe397Leu, Thr448Ala |  |
| Tri-121 | India | Skin, groin | 2016 | *T. indotineae* |  | **>16** | Thr448Ala |  |
| Tri-122 | India | Skin, groin | 2016 | *T. indotineae* |  | **1** | Wildtype |  |
| Tri-123 | India | Skin, groin | 2016 | *T. indotineae* |  | **>16** | Phe397Leu, Thr448Ala |  |
| Tri-124 | Japan | Skin | 2019 | *T. indotineae* |  | **>16** | Phe397Leu |  |

**Supplementary Table S2: Details of whole genome alignments of isolates sequenced as part of this study. NA = not available.**

| Study number | Sequencing depth (x) | *T. indotineae*  reference genome  GCA_023065905.1 | | | *T. interdigitale*  reference genome  GCA_019359935.1 | *T. mentagrophytes*  reference genome  GCA_003664465.1 |
| --- | --- | --- | --- | --- | --- | --- |
|  |  | Reads Mapped (%) | Genome Coverage (%) | Filtered SNPs (N) | Reads  Mapped (%) | Reads  Mapped (%) |
| Tri-01 | 57 | 97.93 | 99.32 | 185 | N/A | N/A |
| Tri-02 | 44 | 95.33 | 98.58 | 115358 | 97.01 | N/A |
| Tri-03 | 57 | 98.01 | 99.34 | 131 | N/A | N/A |
| Tri-04 | 43 | 93.88 | 98.59 | 109089 | 97.09 | N/A |
| Tri-05 | 47 | 97.77 | 99.32 | 194 | N/A | N/A |
| Tri-06 | 54 | 98.21 | 99.34 | 185 | N/A | N/A |
| Tri-07 | 53 | 98.19 | 99.34 | 106 | N/A | N/A |
| Tri-08 | 47 | 98.22 | 99.33 | 337 | N/A | N/A |
| Tri-09 | 54 | 97.89 | 99.33 | 124 | N/A | N/A |
| Tri-10 | 51 | 97.77 | 99.33 | 346 | N/A | N/A |
| Tri-11 | 50 | 98.27 | 99.33 | 338 | N/A | N/A |
| Tri-12 | 52 | 97.34 | 99.33 | 105 | N/A | N/A |
| Tri-13 | 49 | 97.91 | 99.34 | 53 | N/A | N/A |
| Tri-14 | 49 | 99.09 | 99.34 | 181 | N/A | N/A |
| Tri-15 | 51 | 97.31 | 99.33 | 48 | N/A | N/A |
| Tri-16 | 51 | 97.87 | 99.33 | 53 | N/A | N/A |
| Tri-17 | 51 | 97.71 | 99.33 | 115 | N/A | N/A |
| Tri-18 | 47 | 95.71 | 98.55 | 130617 | N/A | 99.05 |
| Tri-19 | 45 | 97.72 | 99.34 | 48 | N/A | N/A |
| Tri-20 | 36 | 97.65 | 99.33 | 117 | N/A | N/A |
| Tri-21 | 47 | 97.77 | 99.33 | 341 | N/A | N/A |
| Tri-22 | 46 | 98.14 | 99.33 | 123 | N/A | N/A |
| Tri-23 | 50 | 98.29 | 99.34 | 122 | N/A | N/A |
| Tri-24 | 50 | 97.54 | 99.33 | 134 | N/A | N/A |
| Tri-25 | 45 | 97.93 | 99.32 | 135 | N/A | N/A |
| Tri-26 | 51 | 98.17 | 99.33 | 114 | N/A | N/A |
| Tri-27 | 54 | 97.97 | 99.34 | 345 | N/A | N/A |
| Tri-28 | 47 | 97.52 | 99.33 | 52 | N/A | N/A |
| Tri-29 | 55 | 97.97 | 99.33 | 334 | N/A | N/A |
| Tri-30 | 46 | 97.96 | 99.34 | 69 | N/A | N/A |
| Tri-31 | 43 | 97.54 | 99.33 | 114 | N/A | N/A |
| Tri-32 | 42 | 97.90 | 99.32 | 114 | N/A | N/A |
| Tri-33 | 47 | 97.67 | 99.33 | 123 | N/A | N/A |
| Tri-34 | 43 | 97.91 | 99.32 | 106 | N/A | N/A |
| Tri-35 | 39 | 97.70 | 99.32 | 104 | N/A | N/A |
| Tri-36 | 2 | 6.24 | 86.28 | 3379 | 6.65 | 6.82 |
| Tri-37 | 32 | 95.78 | 99.35 | 119 | N/A | N/A |
| Tri-39 | 40 | 96.01 | 98.48 | 114654 | 97.61 | N/A |
| Tri-40 | 43 | 98.22 | 99.32 | 52 | N/A | N/A |
| Tri-41 | 38 | 96.93 | 99.31 | 110 | N/A | N/A |
| Tri-42 | 43 | 98.12 | 99.32 | 117 | N/A | N/A |
| Tri-43 | 40 | 97.62 | 99.33 | 136 | N/A | N/A |
| Tri-44 | 43 | 97.45 | 99.33 | 78 | N/A | N/A |
| Tri-46 | 38 | 97.58 | 99.32 | 111 | N/A | N/A |
| Tri-47 | 44 | 97.77 | 99.33 | 114 | N/A | N/A |
| Tri-48 | 43 | 97.65 | 99.34 | 113 | N/A | N/A |
| Tri-49 | 45 | 98.10 | 99.33 | 48 | N/A | N/A |
| Tri-52 | 34 | 97.33 | 99.31 | 184 | N/A | N/A |
| Tri-53 | 51 | 98.17 | 99.32 | 124 | N/A | N/A |
| Tri-54 | 59 | 97.91 | 99.34 | 341 | N/A | N/A |
| Tri-56 | 37 | 98.26 | 99.31 | 59 | N/A | N/A |
| Tri-57 | 39 | 98.01 | 99.33 | 41 | N/A | N/A |
| Tri-58 | 37 | 97.91 | 99.29 | 121 | N/A | N/A |
| Tri-59 | 34 | 97.39 | 99.31 | 50 | N/A | N/A |
| Tri-61 | 36 | 96.65 | 99.31 | 119 | N/A | N/A |
| Tri-63 | 35 | 98.87 | 99.31 | 120 | N/A | N/A |
| Tri-64 | 39 | 97.12 | 99.33 | 115 | N/A | N/A |
| Tri-66 | 45 | 97.65 | 99.33 | 113 | N/A | N/A |
| Tri-67 | 57 | 95.49 | 99.35 | 118 | N/A | N/A |
| Tri-68 | 67 | 98.15 | 99.34 | 113 | N/A | N/A |
| Tri-69 | 40 | 98.41 | 99.32 | 44 | N/A | N/A |
| Tri-70 | 69 | 97.85 | 99.35 | 338 | N/A | N/A |
| Tri-71 | 61 | 98.02 | 99.35 | 51 | N/A | N/A |
| Tri-73 | 51 | 96.01 | 99.34 | 51 | N/A | N/A |
| Tri-74 | 53 | 98.52 | 99.33 | 49 | N/A | N/A |
| Tri-77 | 64 | 98.78 | 99.34 | 122 | N/A | N/A |
| Tri-78 | 63 | 98.54 | 99.35 | 52 | N/A | N/A |
| Tri-80 | 37 | 95.95 | 98.49 | 127393 | N/A | 99.54 |
| Tri-84 | 54 | 96.70 | 98.54 | 130519 | N/A | 99.24 |
| Tri-85 | 51 | 96.05 | 98.55 | 130548 | N/A | 99.19 |
| Tri-93 | 0.25 | 5.51 | 2.75 | 3 | 6.66 | 7.30 |
| Tri-94 | 162 | 92.83 | 98.71 | 118382 | 94.86 | N/A |
| Tri-95 | 0.024 | 3.90 | 0.88 | 0 | 5.76 | 6.23 |
| Tri-96 | 145 | 96.24 | 99.39 | 149 | N/A | N/A |
| Tri-97 | 0.057 | 4.50 | 1.43 | 0 | 5.70 | 5.94 |
| Tri-98 | 187 | 95.01 | 99.39 | 60 | N/A | N/A |
| Tri-99 | 167 | 94.77 | 99.39 | 43 | N/A | N/A |
| Tri-100 | 128 | 95.85 | 99.39 | 120 | N/A | N/A |
| Tri-101 | 122 | 94.51 | 99.38 | 127 | N/A | N/A |
| Tri-102 | 137 | 95.34 | 99.38 | 68 | N/A | N/A |
| Tri-103 | 54 | 30.14 | 99.37 | 112 | N/A | N/A |
| Tri-104 | 179 | 96.39 | 99.39 | 52 | N/A | N/A |
| Tri-105 | 124 | 96.75 | 99.35 | 46 | N/A | N/A |
| Tri-106 | 121 | 93.99 | 99.41 | 146 | N/A | N/A |
| Tri-107 | 89 | 96.50 | 99.37 | 106 | N/A | N/A |
| Tri-108 | 132 | 95.54 | 99.42 | 111 | N/A | N/A |
| Tri-109 | 147 | 91.39 | 99.42 | 108 | N/A | N/A |
| Tri-110 | 173 | 97.10 | 99.35 | 141 | N/A | N/A |
| Tri-111 | 55 | 85.39 | 99.40 | 25 | N/A | N/A |
| Tri-112 | 38 | 85.33 | 99.40 | 39 | N/A | N/A |
| Tri-113 | 32 | 33.66 | 99.34 | 38 | N/A | N/A |
| Tri-114 | 92 | 94.53 | 99.37 | 136 | N/A | N/A |
| Tri-115 | 118 | 95.19 | 99.35 | 108 | N/A | N/A |
| Tri-116 | 112 | 96.55 | 99.36 | 40 | N/A | N/A |
| Tri-117 | 96 | 89.17 | 99.35 | 29 | N/A | N/A |
| Tri-118 | 86 | 97.56 | 99.37 | 42 | N/A | N/A |
| Tri-119 | 132 | 95.36 | 99.33 | 118 | N/A | N/A |
| Tri-120 | 145 | 92.54 | 99.36 | 136 | N/A | N/A |
| Tri-121 | 118 | 92.47 | 99.35 | 100 | N/A | N/A |
| Tri-122 | 36 | 85.42 | 99.39 | 20 | N/A | N/A |
| Tri-123 | 139 | 94.15 | 99.42 | 131 | N/A | N/A |
| Tri-124 | 129 | 98.14 | 99.40 | 116 | N/A | N/A |

**Supplementary Table S3. Average (mean) pairwise SNP difference between *T. indotineae* isolates in each country.** N/A = not available as *n* = 1.

|  | **UK** | **Canada** | **France** | **India** | **Japan** | **Ireland** |
| --- | --- | --- | --- | --- | --- | --- |
| **UK** | 182 | 195 | 157 | 200 | 375 | 128 |
| **Canada** |  | 177 | 167 | 163 | 127 | 118 |
| **France** |  |  | 152 | 122 | 122 | 119 |
| **India** |  |  |  | 88 | 115 | 105 |
| **Japan** |  |  |  |  | N/A | 59 |
| **Ireland** |  |  |  |  |  | N/A |

**Supplementary Table S4. Pairwise *F*_ST_ (Weir & Cockerham) to assess between country diversity.** Values are rounded to 3 d.p.

|  | **Canada** | **France** | **India** | **Ireland** | **Japan** |
| --- | --- | --- | --- | --- | --- |
| **France** | 0.006 |  |  |  |  |
| **India** | 0.038 | 0.013 |  |  |  |
| **Ireland** | -0.123 | -0.084 | 0.008 |  |  |
| **Japan** | -0.105 | -0.079 | -0.018 | 0.000 |  |
| **UK** | -0.001 | -0.007 | 0.024 | -0.101 | -0.097 |

**Supplementary Figure S1: Microreact project of Trichophyton species complex showing a) species identification and b) *SQLE* polymorphisms**

**Supplementary Figure S2: Maximum likelihood phylogeny of all *Trichophyton* isolates (*n* = 99) included in this study, with bootstrap support over 500 replicates.** Branch length represent the average number of SNPs, and branch labels indicate the bootstrap support.

**Supplementary Figure S3: a)** Principle components analysis for all *Trichophyton* isolates included in this study, coloured by confirmed species (*T. mentagrophytes*, *T. interdigitale*, or *T. indotineae*), including the *T. indotineae* type strain. b) Principle components analysis for all *Trichophyton* isolates included in this country coloured by country of isolation.

**Supplementary Figure S4:** a) Within-country diversity assessed by mean pairwise SNP distances for all countries with greater than one isolate available. b) Venn diagram of unique and common SNPs to all isolates sampled in Canada, France, UK and India. c) Estimating between-country divergence with *F*_ST_ (Weir & Cockerham).

**Supplementary Figure S5:** Maximum likelihood phylogeny of *T. indotineae* isolates (*n* = 91) within this study, including the type strain CBS 146623 from Japan, with bootstrap support over 500 replicates. Branch lengths represent the average number of SNPs, and branch labels indicate bootstrap support.

**Supplementary Figure S6: Terbinafine MIC distribution amongst *T. indotineae* isolates (*n* = 90) included in this study, excluding the type strain.** Terbinafine MIC ≥1 mg/L is considered non-wildtype and resistant, and therefore shown in orange.
